# Determination of biophysical parameters for enhancement of human osteoblast proliferation by mechanical vibration in the infrasonic frequency range

**DOI:** 10.1101/448225

**Authors:** Rosenberg Nahum, Halevi Politch Jacob, Rosenberg Orit, Abramovich Haim

**Author notes:** Corresponding author: Rosenberg Nahum MD, FRCS(England), Dept. of Orthopedic Surgery, Rambam Health Care Campus, P.O.Box 9602, Haifa 31096, Israel.

## Abstract

Experimental methods for studying an enhancement of osteoblast proliferation in vitro provide tools for the research of biochemical processes involved in bone turnover in vivo. Some of the current methods used for this purpose are based on the ability of the osteoblasts to enhance proliferation by mechanical stimulation. We describe an experimental approach of biomechanical stimulation of cultured human osteoblast-like cells by vibration. This method is based on the specially designed controlled vibration setup that consists of an electric actuator, with horizontally mounted well plate containing cell cultures. Previously this method found to be effective to enhance cell proliferation, but the exact mechanical parameters of effective vibration were elusive. The current low friction system for mechanical stimulation of osteoblast-like cells in vitro provides recording of narrow range mechanical parameters in the infrasonic spectrum.

We exposed human osteoblast-like cells in explant monolayer culture to mechanical vibration in the 10-70Hz range of frequencies and found that 50-70 Hz of vibration frequency is optimal for inducing osteoblast proliferation that was deduced from interrelation between unchanged cell number in culture samples with significant decrease in cell death rate (decreased LDH activity in culture media, p<.05) and with parallel decrease of their maturation level (p<.01).

In this report we determined the optimal mechanical parameters and excitation protocol for induction of osteoblast proliferation in vitro by using a tunable and versatile mechanical platform, which can be used in the research of cell mechanotransduction.

## Background

Mechanical loading increases bone mass by activating osteoblast metabolic activity and proliferation via cellular biomechanical pathways. These pathways are studied in vitro in models of cellular mechanical stimulation. The widely used experimental methods for this purpose utilize stretching of cells adherent to elastic membranes or implementation of controlled external fluid flow to cell cultures [1, 2]. In both, the mechanism of cellular activation is due to shearing forces and subsequential cellular deformation, which causes cytoskeletal activation, mainly involving the microtubular and microfilament components [3, 4]. These experimental methods require sophisticated hardware with essential special handling and complex experimental protocols, but usually unable to produce uniform stretching forces on all the cells in culture, and commonly restricted to the mechanical loading frequencies up to 5Hz [5]. Previously a more versatile experimental approach has been proposed which is highly reproducible and with much wider range of mechanical frequencies that can be applied on the cultured osteoblasts in monolayer. This method is based on application of controlled vibration force in a defined range of mechanical parameters [6].

In this model, the cellular deformation is caused by shearing forces, applied by extra-an intracellular fluid flow, which is induced by the one-dimensional accelerated vibration movement of cells, adherent to a plastic surface. Since the cells in this model are in the same environmental condition, they exposed to the same magnitude of force during the vibration force application on a culture well-plate. Therefore, large numbers of cells can be studied in the same mechanical conditions, which are determined by the dimensions of the well plate and its weight, its surface biophysical properties, volume of the culture media in the wells and by a profile of the force generated by a vibration actuator. Additionally, this experimental setup can be used for application of vibration in higher frequencies, i.e. in the infrasonic range (10-70 Hz).

There are several previous reports that human osteoblast in vitro can be stimulated in the infrasonic range of vibration frequencies [3, 6, 7], probably by mimicking the natural influence of resting muscle on an attached bone, as it is seen on the vibromyogram [8] following muscle’s spontaneous periodic contraction in this range of frequencies. These reports are also supported by the in vivo studies that showed an enhancing effect of vibration in this range of frequencies on the bone mass buildup [9].

It has been shown that different sub ranges of effective vibration parameters for osteoblast proliferation and metabolic activity stimulation exist and not overlapping [6]. The main difficulty that evolved in that model was to keep a constant pattern of vibration parameters and to record these values directly from the moving supporting well plate, due to mechanical instability of the electromagnetic actuator and the difficulty of the fine-tuning of the mechanical parameters in this range of frequencies. The source of the variability of vibration signal was the surface friction of the moving parts, i.e. in that model it was impossible to effectively eliminate or reduce the additional vibration frequencies, in higher or lower range, which were produced by the vibration actuator and by the well plate movement. Because the optimal range of vibration frequencies for osteoblast activation is narrow, the delivery of precise and stable external vibration signal is crucial. Additionally, to claim that infrasonic range of frequencies has an enhancing effect on cultured osteoblast, a “pure” sinusoidal vibration force pattern, or very close to it, should be applied. Currently, a report of such experimental setup does not exist. Therefore, to determine the role of mechanical excitation of osteoblast in the infrasonic range of frequencies we developed an experimental model for human osteoblast stimulation by external vibration using a low friction system for delivery of vibration force to the culture well plate. This system provides a closer to the “pure” sinusoidal vibration force that can be effectively controlled and measured.

We hypothesize that by using the designed precise method of vibration force application we could provide a reliable evidence that human osteoblast is stimulated in the infrasonic range of mechanical vibration and provide a more precise data on these effective mechanical parameters.

## Methods

### Cell culture protocol

The experiments were performed on cultured osteoblast-like cells originated from disposable cancellous bone samples taken from distal femurs of human donors during osteoarthritic knee arthroplasties. The method was approved by the institutional Ethical Committee. Bone chips of cancellous bone, 2 – 3 grams in total, were incubated in DMEM with heat-inactivated fetal calf serum (10%), 20mM HEPES buffer, 2mM L-Glutamine, 100μM Ascorbate-2-Phosphate, 10nM Dexametasone, 50 U/ml Penicillin, 150μ/ml Streptomicin at 37°C in humidified atmospheric environment of 95% air with 5% CO_2_ (v:v) for 20 - 30 days. Human osteoblast-like cells grow out from the chips as a primary cell explants cultures adherent to the plastic tissue culture plates. The human bone cell cultures obtained by this method have been shown previously to express osteoblast-like characteristics such as polygonal multipolar morphology, expression and activity of the enzyme alkaline phosphatase, positive Von Kossa staining, synthesis of osteopontin, synthesis of a collagen-rich extracellular matrix with predominantly type I collagen and small amounts of collagen type III and V, as well as noncollagenous proteins such as sialoprotein (BSP) and osteocalcin. Additionally, these cells demonstrate matrix mineralization *in vitro* and bone formation *in vivo* [10, 11, 12]. The cells were passage into 24 well plates. Each well will be seeded with 2X10^4^ cells.

### Mechanical system configuration

Plastic 24 well-plate with the cultured osteoblast-like cells (weight 120 grams including the culture media), was mounted on a stage that moves on a low friction support (Figure 1). The mechanical force was applied to the well plate by an electromagnetic actuator which is horizontally connected to the moving stage. A piezoelectric accelerometer (accelerometer sensitivity = 100mV/g,) is attached to the well plate in the horizontal direction of movement. The relation between the output voltage from a piezoelectric accelerometer and the acceleration is linear throughout frequencies of vibration of 10-60Hz and at the displacement amplitude of up to 30μm.

**Figure.1.**
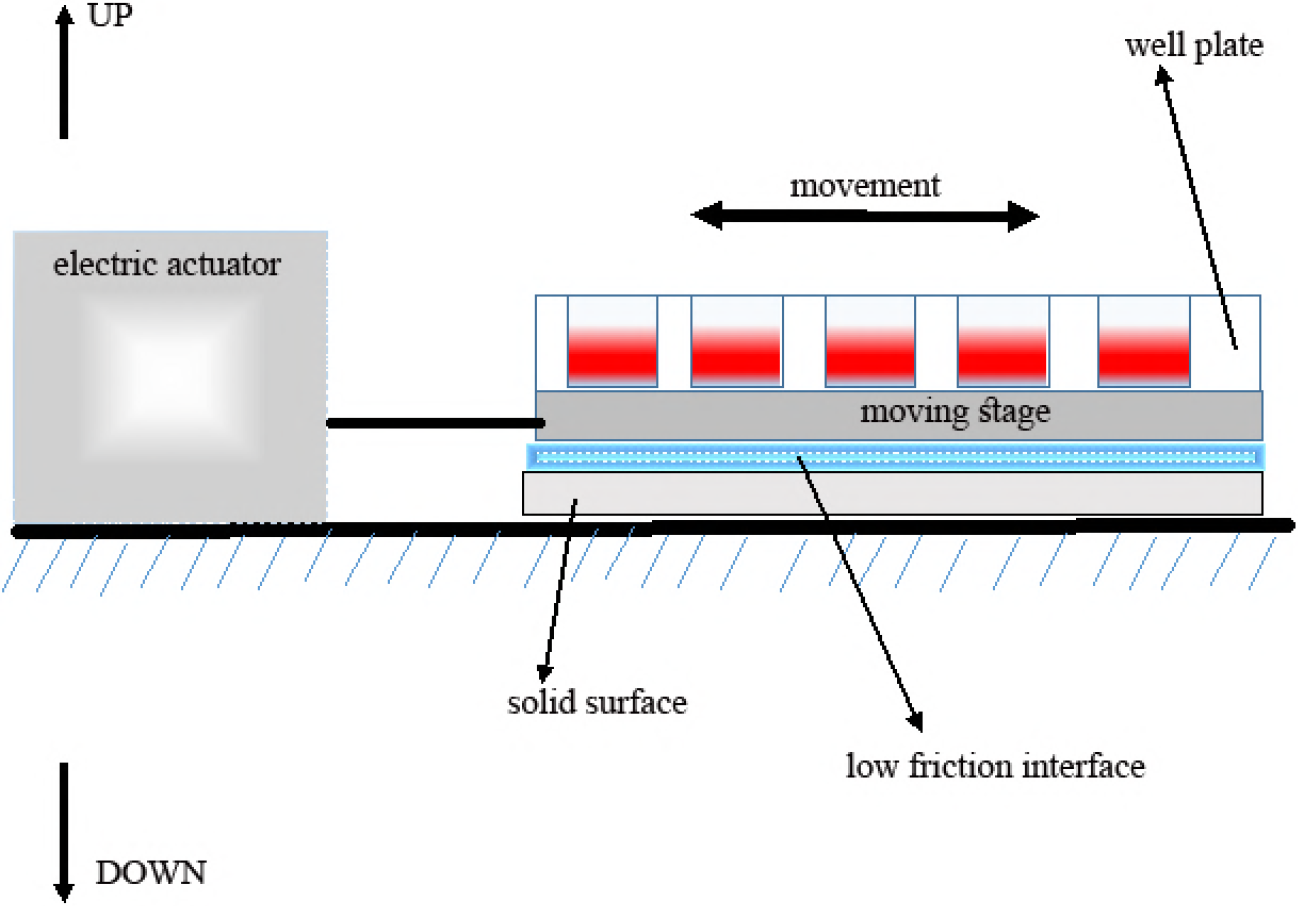
Schematic representation of the experimental setup for mechanical stimulation by vibration movement. Plastic 24 well-plate with culture samples of cells adherent to the plastic surface and covered with culture media is firmly attached to the moving stage on a low friction interface. The vibration movement is controlled by the electric actuator. The movement parameters are recorded by the accelerometer, which is attached to the moving stage.

This system produces a vibration movement, which is close to sinusoidal, with displacement amplitudes in the range of 20 to 30 μm and a near sinusoidal vibration frequency of 10 and 60 Hz (Figure 2).

**Figure.2.**
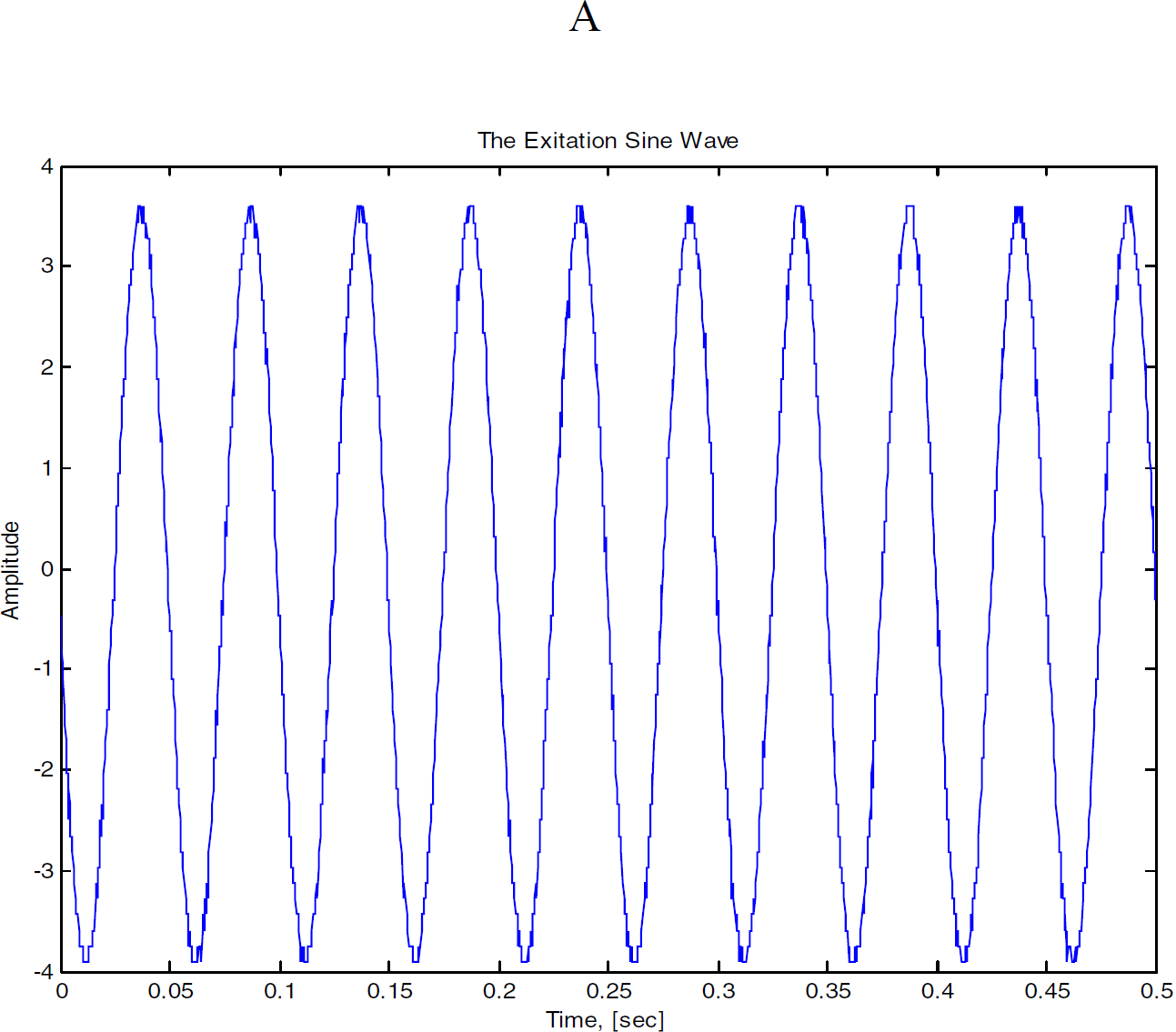
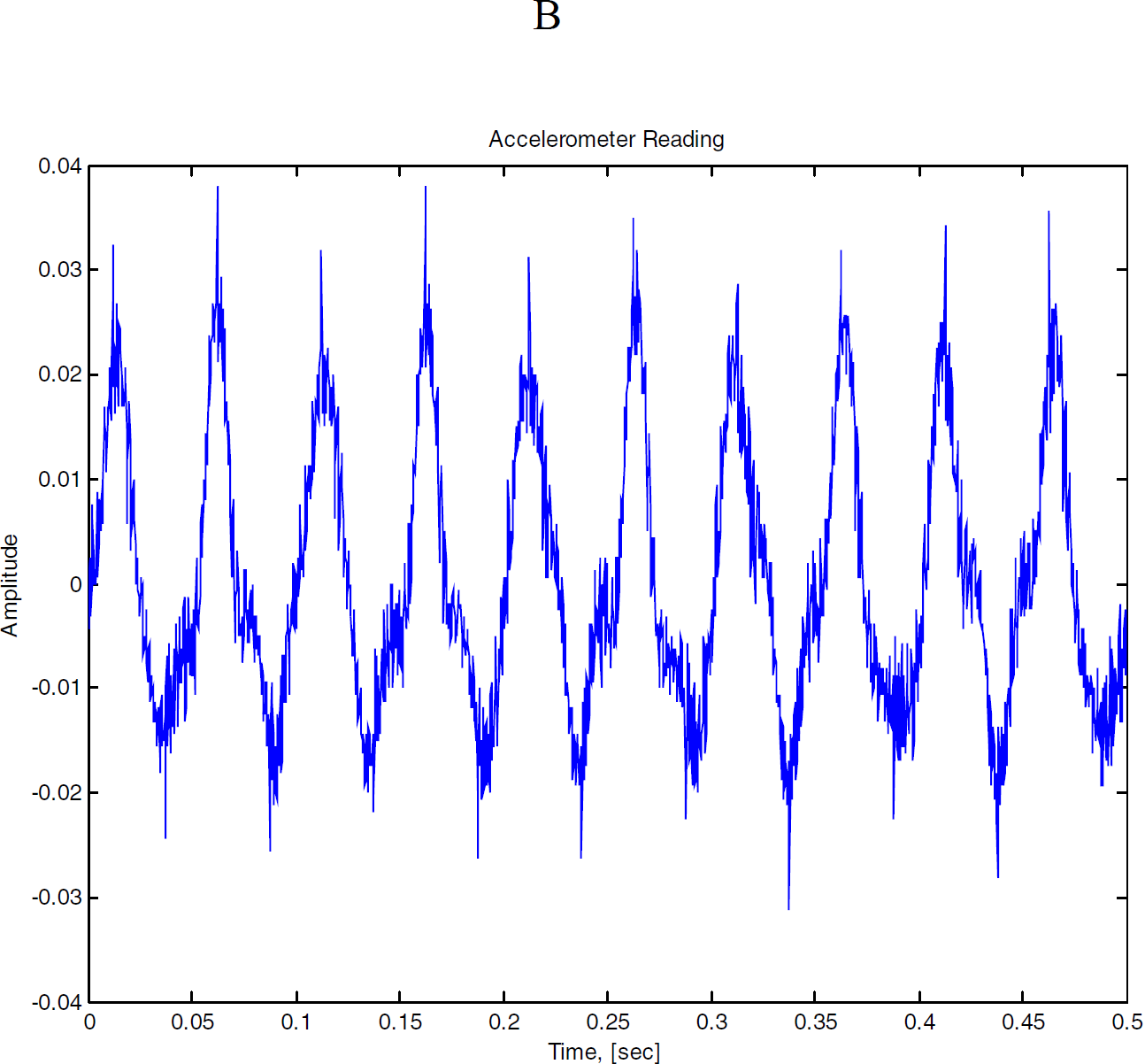
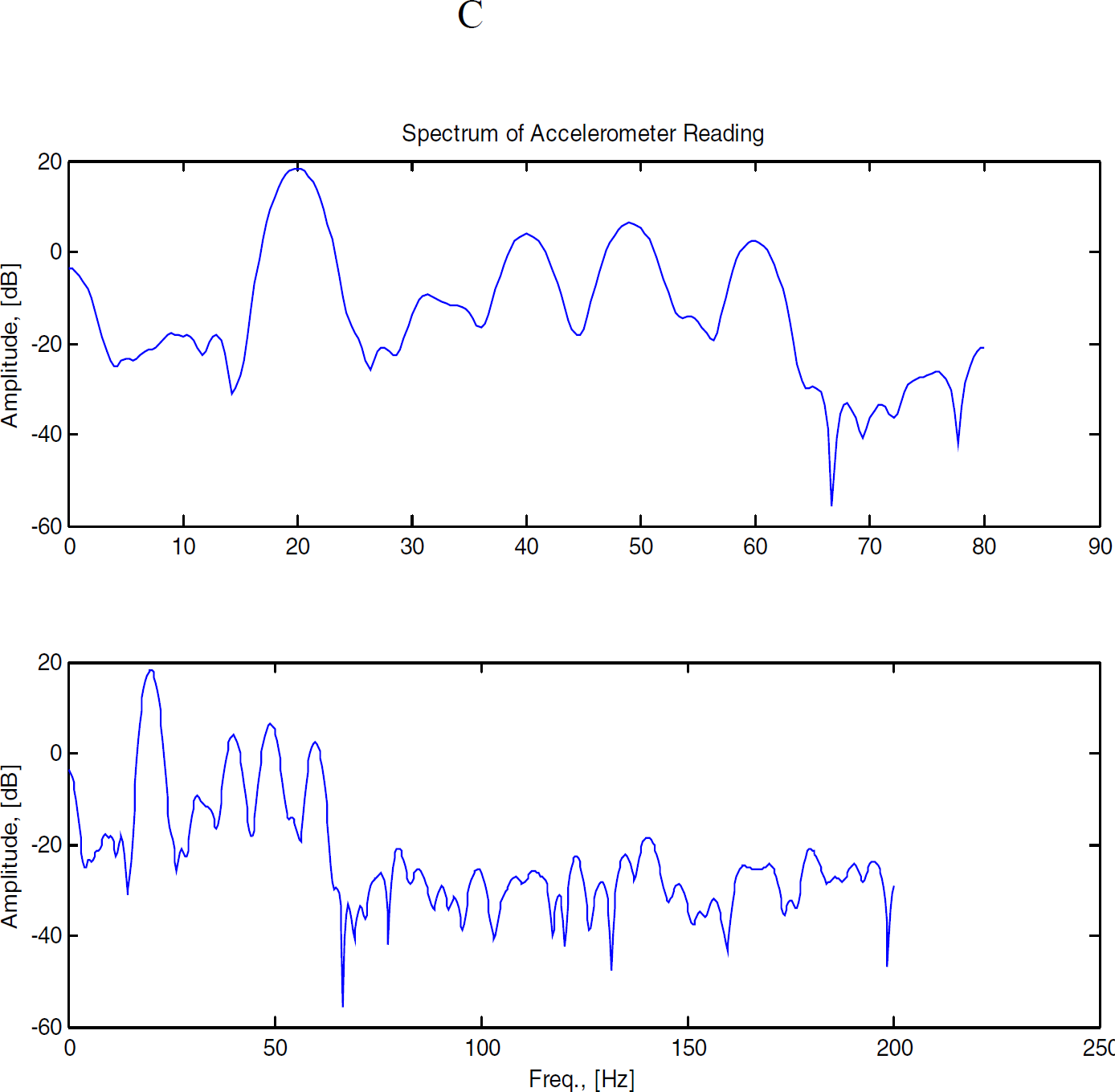
Example of the excitation profile of 20 Hz excitation by the actuator (A) and the measurements readings by the accelerometer attached to the moving stage (B), which represents the actual mechanical force applied to cells adherent to the moving surface and appears close to the excitation signal from the actuator (amplitude – V*10^-1^). The spectra of the vibration frequencies, which were applied to the cells (C) shows that 70% of the frequencies, up to 70 Hz frequency, are in the range of 10-30 Hz.

By the readings from the piezoelectric accelerometer we could define the dominant frequencies and recognize existence of additional harmonic frequencies (Figure 2). Through a double integration, the displacement of the well plate movement can be measured.

The spectra of the vibration frequencies applied to the cells show that when the system is tuned to excitation of 20 Hz vibration, 70% of the frequencies (up to 70 Hz frequency) are in the range of 10-30 Hz, and when the system is tuned to excitation of 60 Hz vibration, 98% of the frequencies (up to 70 Hz frequency) are in the range of 50-70 Hz. Therefore, by the main excitation from the actuator a specific narrow range band of frequencies that are applied directly to the cultured cells was generated.

### Mechanical stimulation protocol of the cultured osteoblast-like cells

Following first passage of cells, replicates of equal number of cultured cells were submitted to vibration conditions of 20Hz range or 60 Hz range. Duration and timing of exposure of cultured replicates to vibration were according to previously found optimal protocols [3, 6], i.e. four repetitions of two minutes exposure at 24 hours intervals. The culture samples which kept in static conditions, without exposure to vibration protocols, were used as controls. Twelve cell culture samples for each experimental condition were studied.

Number of cells in each culture sample was estimated by direct counting in low power microscopic field (by using ImageJ computerized picture processing). Mean value of cell number from three different microscopic fields in each culture sample, after exposure to mechanical stimulation and in control samples that were kept at static condition, was recorded. We preferred this method of cell number evaluation because it avoids interfering with the culture conditions.

### Cell death and proliferation estimation

LDH activity in the culture media, a marker of cell death due to LDH leakage via damaged cell membranes, was measured in the samples exposed to mechanical vibration and in the static controls. Briefly, LDH activity in culture media samples was determined by 340 nm wavelength spectrophotometry of the reduced nicotineamide adenine dinucleotide (NAD), which is directly proportional to LDH activity [13, 14]. Accordingly, the cell proliferation comparison among the studied samples in different conditions, i.e. exposed to 60Hz, 20Hz of vibration protocols and static controls, was based on a result of the deduction of cell death change from the total cell number in the culture samples.

### Cellular maturation estimation

Cellular alkaline phosphatase activity, which is a marker of osteoblast maturation and inversely related to osteoblast proliferation. The cellular alkaline phosphates activity was measured by spectrophotometry [15]. Briefly, following microscopic cell counting of each culture sample, the medium was removed and cells, adherent to the plastic surface, were washed by PBS and lysed in 0.1% Triton X-100 by 3 cycles of freezing at -20°C and thawing at 20°C. Alkaline phosphatase activity was determined in the lysed cell culture samples, after incubation with P-nitrophenyl phosphate substrate, by 410 nm wavelength spectrophotometry

### Data analysis

The comparison of two parameters was done by the t test and the comparison of three parameters by ANOVA, and subsequentially by the Dunnett’s post hoc test (following verification of normal distribution of results and with p value below 0.05). P value of less than 0.05 was considered as an evidence of significant difference between the compared variables.

## Results

Following the 60Hz vibration protocol the number of cells significantly increased in comparison to cells kept in control static condition (average 68 ± 4 SEM cells/microscopic field vs. average 55 ± 4 SEM cells/microscopic field respectively, n=12, one-way ANOVA p<0.05, Fig. 3). No significant change in number of cells exposed to 20Hz protocol in comparison to static controls was observed (average 64 ± 3 SEM cells/microscopic field vs. average 55 ± 4 SEM cells/microscopic field respectively, n=12, one way ANOVA p>0.05, Fig.3), There was no difference in the media LDH activity between the static controls and cultures exposed to either 20Hz or 60Hz vibration protocols (average 193 ± 12 SEM U/L vs. average 163 ± 9 SEM U/L and average 167 ± 8 SEM U/L respectively, n=12, one-way ANOVA, p>0.05).

**Figure.3.**
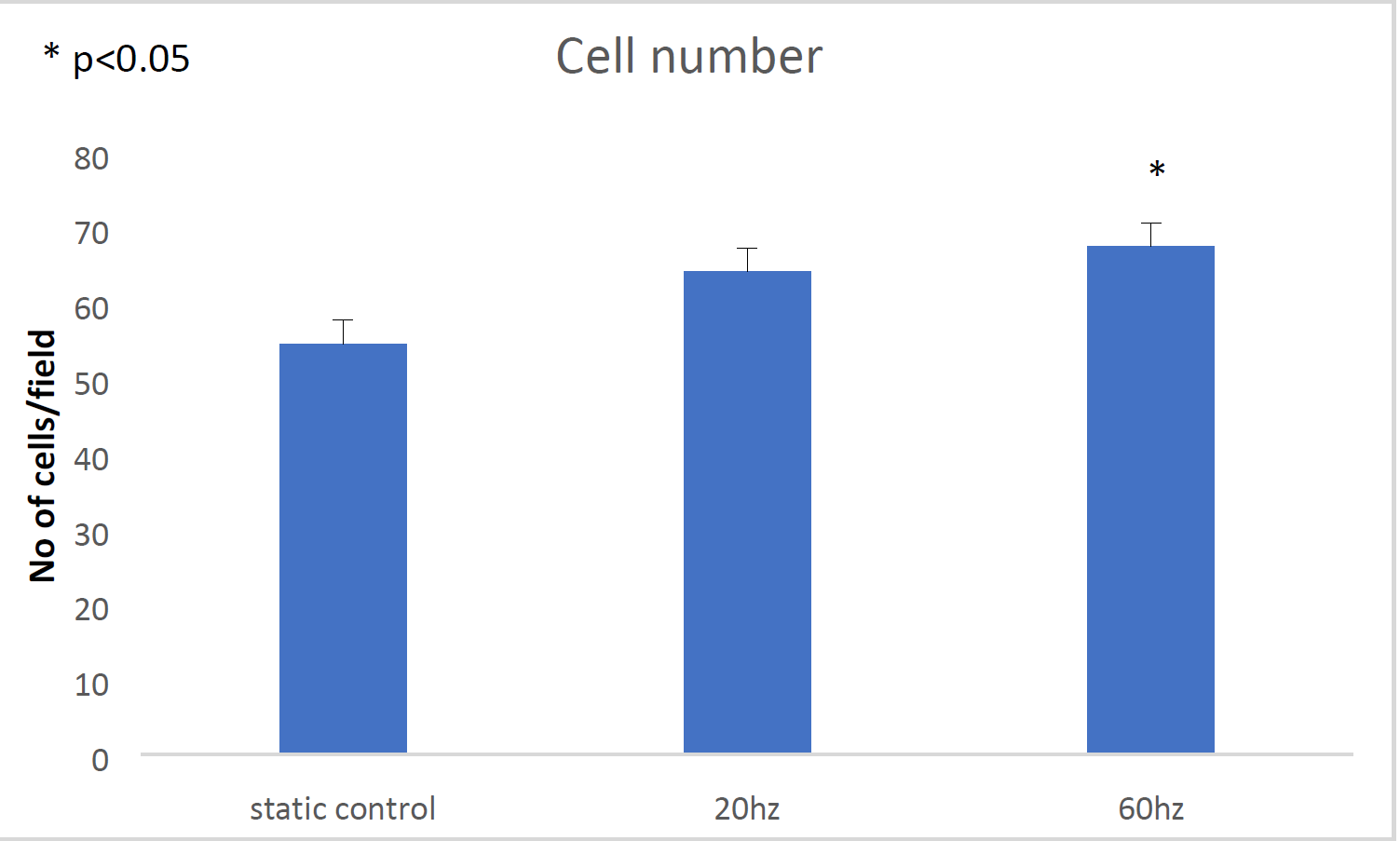
Number of cells raised significantly following 60 Hz vibration stimulation (p<.05, n=12). Average values in culture samples with SEM (vertical bars) are presented.

We found a significant decrease in cellular alkaline phosphatase activity following 60Hz vibration protocol in comparison to static controls (average 0.559 ± 0.041 SOM U/L/cell in microscopic field vs. average 0.794 ± 0. 0571 SEM U/L/ cell in microscopic field respectively, n=12, t test, p<0.01, Fig. 4).

**Figure.4.**
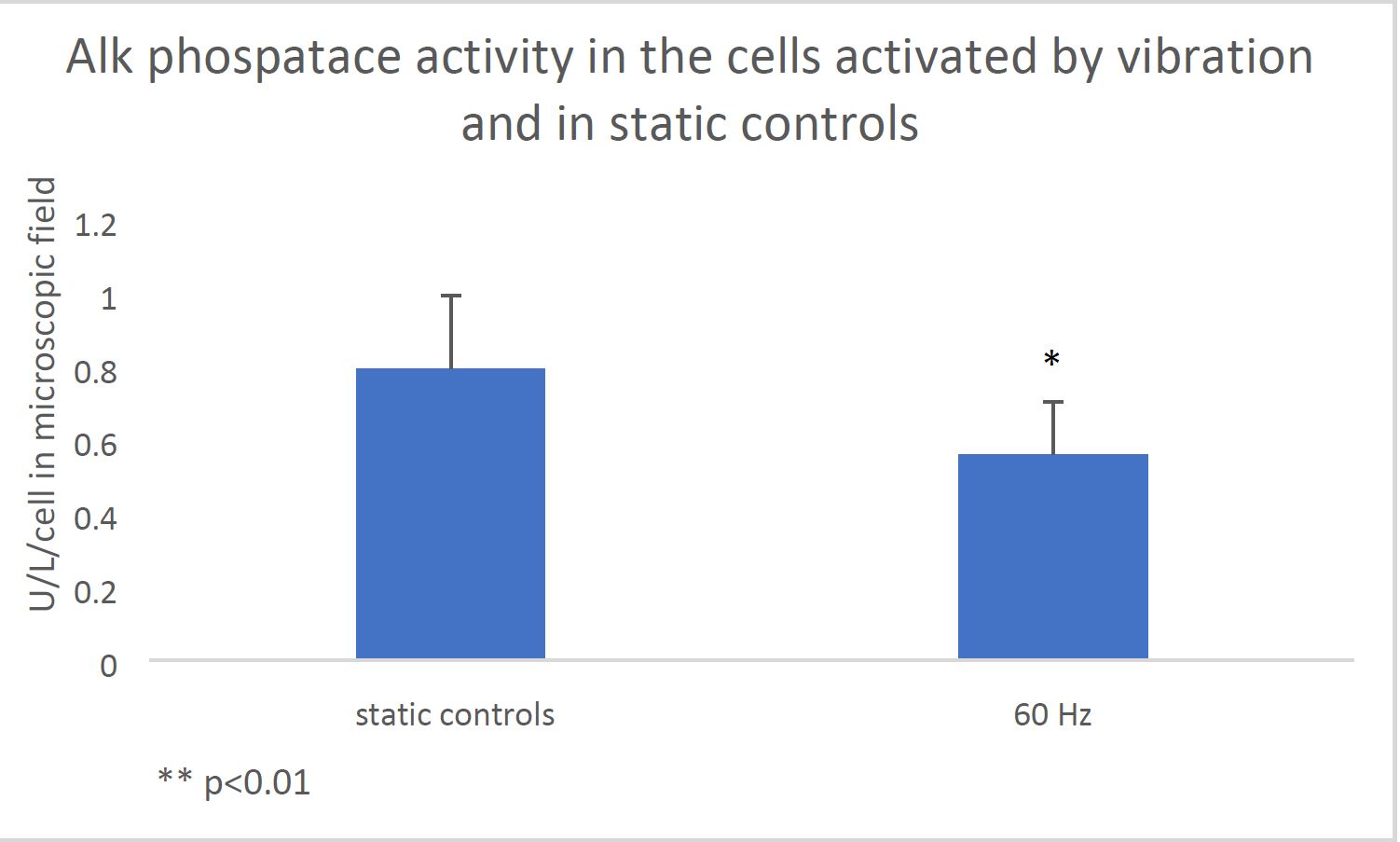
After the 60 HZ vibration, the cells’ maturation state decreased-significantly lower alkaline phosphatase activity in the mechanically activated cells was found (p<0.01, n=12). Average values in culture samples with SEM (vertical bars) are presented.

## Discussion

In this study, we determined the precise range of frequencies, in the infrasonic spectrum of 50-70 Hz (when excitation by 60Hz provided by the actuator) that is specifically effective in enhancing human osteoblast proliferation in vitro. This conclusion is following the deduction of cell death rate from cell numbers per sample in different mechanical protocols that we studied, i.e. when 60 Hz excitation of cells didn’t cause a significant change in cell number in comparison to controls, but did cause a significant decrease in cell death (lower LDH activity in culture media) the conclusion is that this type of mechanical stimulation did cause an enhancement in cell proliferation. This phenomenon wasn’t apparent following 20Hz excitation. Furthermore, this conclusion is independently supported by the evidence of the lower maturation state of the cells exposed to the 50-70Hz mechanical vibration protocol. We preferred this double approach of independent assays to increase the confidence in the results. Indeed, the results show that following exposure to the upper spectrum of the infrasonic vibration (around 60 Hz) the cells react by increased proliferation and stay in lower differentiation state, as expected. This effect doesn’t happen in the lower range of the infrasonic spectrum (cellular excitation by 20Hz that provide 10-30 Hz of vibration spectrum), i.e. by this excitation protocol neither the cell number nor LDH activity in culture media were significantly different from the cells kept in control static environment, This fact supports our basic hypothesis that in the infrasonic range of mechanical cellular excitation, only a specific band of frequencies of vibration elicits a proliferation response.

This observation is different from the previous report on a more pronounced proliferation response by 20Hz excitation [6]. We attribute this controversy to a less precise experimental setup in the previous report, when the actual mechanical parameters of the moving surface were unavailable, therefore the actual excitation spectrum is unknown and might had contained additional mechanical frequencies that were generated by the actuator. One of the main purposes of the present study was to find the narrow range of mechanical frequency, which causes the highest rate of cell proliferation.

Therefore, according to the study hypothesis, we describe a tunable mechanical platform for the research of mechanotransduction in osteoblasts with determination of the optimal mechanical frequencies for induction of cell proliferation. Naturally, other cellular functions, related to mechanical stimulation, can be studied by the utilizing this versatile method.

We didn’t attempt to determine the specific mechanotransduction biomechanical pathways that are evoked by this mechanical excitation method, but rather provided the basic optimal biomechanical data for the further mechanotransduction research by using the presented experimental setup that generates a uniform force transfer to a large number of cells in culture.

## References

1 Schaffer JR, Rizen ML, Italien GJ, Benbrahim A, Megerman J, Gerstenfeld LC, Gray ML. Device for application of a dynamic biaxially uniform and isotropic strain to a flexible cell culture membrane. J Orthop Res 1994; 12: 709–719

2 Soejima K, Klein-Nulend J, Semeins CM, Burger EH. Calvarial and long bone cells derived from adult mice respond similarly to pulsating fluid flow with rapid nitric oxide production. Calcif Tissue Int 1999; 64: S113

3 Rosenberg N. The role of the cytoskeleton in mechanotransduction in human osteoblast-like cells. Human Exp Toxicol 2003; 22: 271–274.

4 Toma CD, Ashkar S, Gray ML, Schaffer JL, Gerstenfeld LC. Signal transduction of mechanical stimuli is dependent on microfilament integrity: Identification of Osteopontin as mechanically induced gene in osteoblasts. J Bone Min Res 1997; 12: 1626–1636

5 Steward AJ, Cole JH, Ligler FS, Loboa EG. Mechanical and vascular cues synergistically enhance osteogenesis in human mesenchymal stem cells. Tiss Eng 2016; 22; 997–1005

6 Rosenberg N, Levy M, Francis M. Experimental model for stimulation of cultured human osteoblast-like cells by high frequency vibration. Cytotechnology 2002; 39: 125–130

7 Wang L, Hsu H, Xian J. Vibration analysis of a single osteoblast in vitro using the finite element method. Int Conf Biomed Biol Eng 2016; 380–385.

8 Nigg BM. Acceleration. In: Biomechanics of the musculo-skeletal system. Second Edition Nigg BM & Herzog W (eds), Chichester: John Wiley & Sons, 1998: 300–301

9 Rubin C, Li C, Sun Y, Fritton C, McLeod K. Non-invasive stimulation of trabecular bone formation via low magnitude, high frequency strain. Trans ORS 1995; 20: 548

10 Rosenberg N, Soudry M, Rosenberg O, Blumenfeld I, Blumenfeld Z. The role of activin A in the human osteoblast cell cycle: A preliminary experimental in vitro study. Exp. Clin. Endocrin Diab 2010; 118; 708–712

11 Gundle R, Stewart K, Screen J, Beresford JN. Isolation and culture of human bone-derived cells. In: Marrow stromal cell culture. Beresford N & Owen ME (eds), Cambridge University Press 1998, Cambridge, UK: pp 43–66

12 Yamanouchi K, Satomura K, Gotoh Y, Kitaoka E, Tobiume S, Kume K, Nagayama M. Bone formation by transplanted human osteoblasts cultured within collagen sponge with dexamethasone in vitro. J Bone Min Res 2001; 16: 857–867

13 Gay RB, Bowers GN Jr. Optimum reaction conditions for human lactate dehydrogenase isoenzymes as they affect total lactate dehydrogenase activity. Clin Chem 1968; 14: 740–753

14 Rosenberg N, Hamoud K, Rosenberg O. Quantitative expression of cell death by LDH activity. IOSR J Pharm Biol Sci 2016; 11: 46–48

15 Bessey OA, Lowry OH, Brock MJ. A method for the rapid determination of alkaline phosphatase with five cubic millimeters of serum. J Biol Chem 1946; 164: 321–329

